# Diseased skin dermal proteomic profiles reflect shared extracellular matrix dysregulation patterns

**DOI:** 10.1101/2024.08.27.609987

**Authors:** Luís Martins, Mariana D. Malta, Sara Chaves, Hugo Osório, Christina Guttmann-Gruber, Thomas Kocher, Alexandra P. Marques

**Affiliations:** 3B’s Research Group, I3Bs-Research Institute on Biomaterials, Biodegradables and Biomimetics, University of Minho, Headquarters of the European Institute of Excellence on Tissue Engineering and Regenerative Medicine, 4805-017 Guimarães, Portugal; ICVS/3B’s-PT Government Associate Laboratory, 4805-017 Guimarães, Portugal; i3S – Instituto de Investigação e Inovação em Saúde, Universidade do Porto, 4200-135 Porto, Portugal; Ipatimup – Instituto de Patologia e Imunologia Molecular da Universidade do Porto, 4200-135 Porto, Portugal; FMUP – Faculdade de Medicina da Universidade do Porto, 4200-319 Porto, Portugal; EB House Austria, Research Program for Molecular Therapy of Genodermatoses, Department of Dermatology and Allergology, University Hospital of the Paracelsus Medical University, Salzburg, Austria

## Abstract

Extracellular matrix (ECM) plays a major role in the maintenance of skin homeostasis and modifications in its structure are commonly linked with skin diseases of different origins. Recent evidence shows that epidermal fibroblasts can be important regulators of epidermal pathology. However, despite recent advances, the intricate mechanisms responsible for the ECM defects in a wide range of skin pathologies are still evasive.

In this work we used Dystrophic Epidermolysis Bullosa, Pemphigus vulgaris and Squamous Cell Carcinoma, as the models for epidermal diseases of distinct etiology, in order to explore potential shared alterations in diseased dermal fibroblasts.

Our proteome analysis revealed that differentially expressed proteins in all diseases are commonly enriched in processes related to supramolecular fiber organization, complex of collagen trimers and actomyosin, however with opposite patterns. Nevertheless, Collagen XII is significantly downregulated in all diseases. Additionally, an algorithmic pipeline predicts that MAPKs and CDKs are major regulators of dermal proteome alterations across all diseases.

Altogether, our results highlight a possible shared mechanism where downregulation of Collagen XII mediates ECM organization disruption leading to diverse disease phenotypes.

## INTRODUCTION

Extracellular matrix (ECM) is the pilar of skin structural stability. Its relationship with multiple cell types in the different cutaneous layers also gives ECM a substantial role in skin homeostasis. Several alterations in ECM proteins have been associated with skin disorders of various etiologies, from heritable and autoimmune to skin cancers [1,2]. Yet, the diversity of ECM-related defects and the small number of people affected by certain cutaneous diseases makes them less appealing for in-depth understanding of their similarities and differences.

Regardless of the etiology, the regulation of disease onset or progression extends beyond the affected skin compartment, likely involving common mediators. Recent data from immune-mediated skin diseases vitiligo and psoriasis has shown that dermal fibroblasts play a critical role in disease pathogenesis by, respectively, defining the pattern of affected melanocytes in the epidermis [3], and acting as signaling intermediaries in basal epidermal cell proliferation [4]. Additionally, transcriptomic comparison of different inflammatory skin diseases demonstrated the presence of shared molecular signatures [5]. Skin blistering diseases such as dystrophic epidermolysis bullosa, caused by mutations in the *COL7A1* gene [6,7], and pemphigus vulgaris (PV), induced by auto-antibodies against DSG1 and 3 in the epidermis [8], are commonly associated with excessive dermal inflammation. Investigation of dermal fibroblasts in DEB has shown increased activity of TGFβ and expression of several ECM proteins that correlate with collagen VII degradation and disease severity [9,10]. Additionally, disruption of collagen VII anchoring fibrils results in reduced strength of epidermal-dermal cohesion, chronic skin fragility, fibrosis and consequently altered ECM organization [6]. Dermal knowledge in PV is scarce. However, an increased inflammatory environment was found in the dermis of PV patients, where accumulation of apoptotic cells was shown to increase the expression of TNF-α by monocytes, further increasing the susceptibility to acantholysis [11], and PV sera was shown to increase the expression of collagen degrading MMP9 in animal models [12]. In what concerns skin cancers, dermal ECM has not been directly linked to the initiation of the tumorigenic process. However, increased expression of enzymes that degrade the basement membrane components laminin and collagen IV were found to be increased in squamous cell carcinoma (SCC), a skin tumor characterized by abnormal keratinocyte proliferation and dermal invasion [13–15], and correlate with tumor invasiveness [16,17]. Moreover, ECM stiffness, resulting from abnormal collagen production and deposition, induces the development of tumor clusters and is linked to increased tumor cells migration and invasiveness [18,19]. Interestingly, aberrant cycles of wound healing resulting in fibrosis have been associated with the higher susceptibility of SCC development in severe DEB patients [7,18].

In line of this evidence, we hypothesized that the identification of common pathogenic disease mechanisms and therapeutic targets could increase the prospects for effective treatments. As such, we investigated the dermal proteome profile of DEB, PV and SCC, all of them affecting the epidermis, in pursuit of shared dermal alterations with potential for therapeutical exploitation. Our proteomic analysis identified a common pattern of disorganization of collagen fibers and actomyosin that could have implications in the pathomechanisms of each disease.

## RESULTS

### Protein expression profile of dermal fibroblasts varies among skin diseases

To get a better understanding of skin pathogenesis and to explore a potential common pattern among dermal fibroblasts of skin diseases we used LC-MS to perform proteome analysis of fibroblasts cultured in over-confluence to promote ECM deposition. We identified a total of 5220 proteins in all disease variants. To visualize the global expression profile we used Uniform Manifold Approximation and Projection (UMAP) [20] which showed that all disease variants and healthy control have different expression patterns (**Figure 1A**). As expected, DDEB, intRDEB and sevRDEB variants presented more similar expression patterns to each other than with the other diseases. The same was observed for PVcut and PVmuc expression patterns.

**Figure 1.**
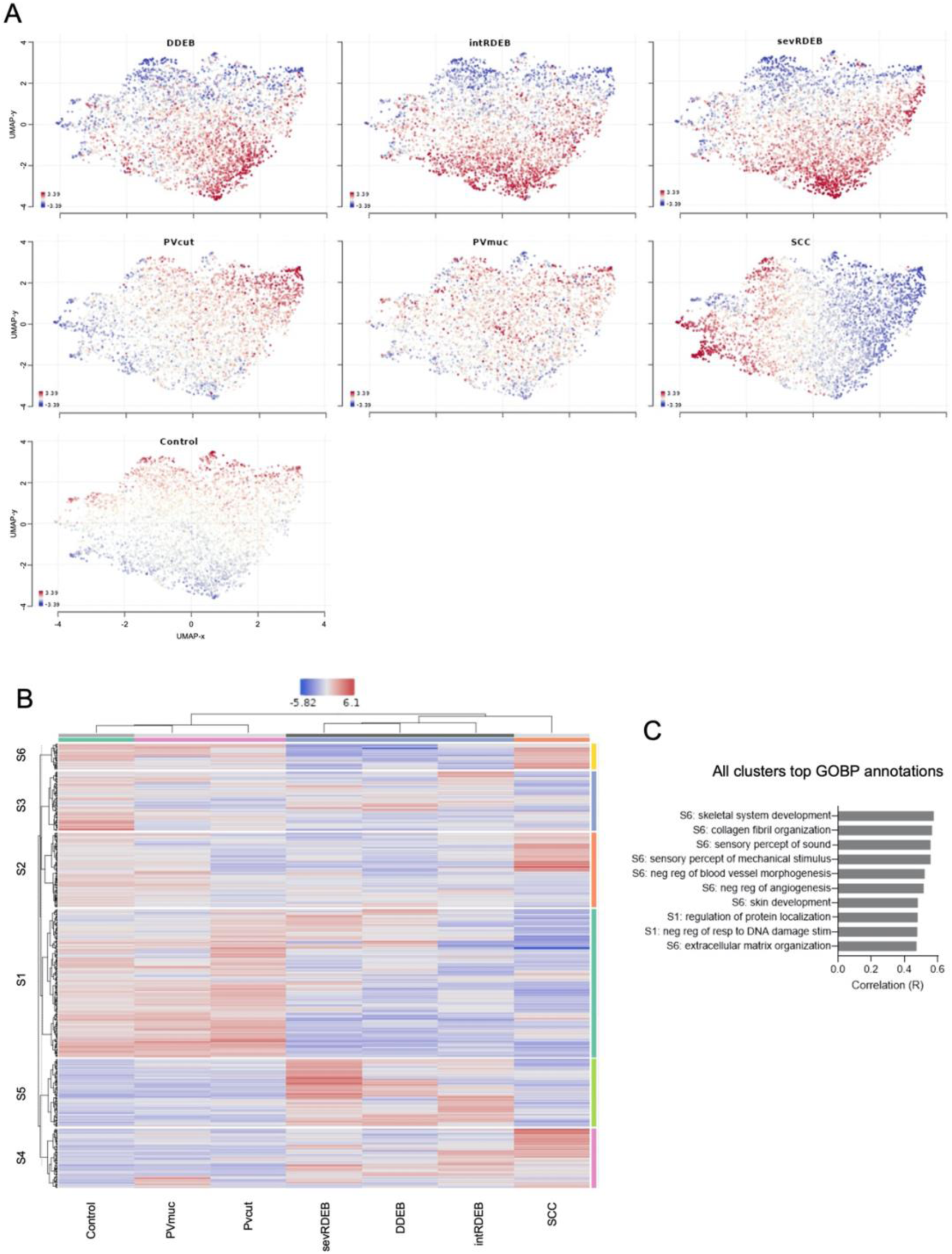
Proteomic profile of dermal fibroblasts of different skin diseases. **A)** Uniform manifold approximation and projection (UMAP) visualization of protein expression coloured by relative normalized log-expression (logCPM). Distance metric between proteins is covariance. Proteins clustered nearby have high covariance; **B)** Heatmap of the 500 top features, defined by standard deviation, showing protein expression sorted by 2-way hierarchical clustering; **C)** Top ranked annotation features (by correlation) using the Gene ontology biological process database (GOBP) for all protein clusters defined in the heatmap.

Next, we performed 2-way hierarchical clustering (**Figure 1B**). The top 500 feature clustered heatmap corroborated that all diseases have different protein expression patterns and clustering supported the pattern of similarities between variants of the same disease. Notably, PVcut and PVmuc were assigned under the same hierarchical branch as the healthy control, indicating fewer alterations despite the significance of the disease. To further comprehend the hierarchical distribution of those top proteins we analyzed the functional cluster annotations using the Gene Ontology Biological Processes (GOBPs) database. Cluster S6 showed the highest correlation scores for GOBP annotations. Nonetheless, the expression pattern of proteins in this cluster was distinct between all diseases or respective variants. Moreover, all clusters top GOBPs correlations were related to skeletal system development and collagen fibril organization. GO terms for skin development and extracellular matrix were also among the top annotations (**Figure 1C**).

### Collagen fibril and actomyosin processes are altered in all diseases but not in the same manner

To further explore the alterations associated with the different diseases we performed differential expression analysis for each disease versus healthy control. We found 1149, 1300, 1333, 107, 276 and 1278 differentially expressed proteins (DEP) respectively in the comparisons DDEB, intRDEB, sevRDEB, PVcut, PVmuc and SCC versus control at a threshold of log2 Fold change (FC) ± 1 and q value < 0.05 (**Figure 2A**). Interestingly, PVcut and PVmuc had the lowest number of DEPs, corroborating the higher degree of similarities with the control shown in the UMAP and clustered heatmap. Pairwise scatterplots comparing the six differential expression contrasts of disease versus healthy control also showed that diseases have closer similarities to variants of the same disease (**Figure S1**).

**Figure 2.**
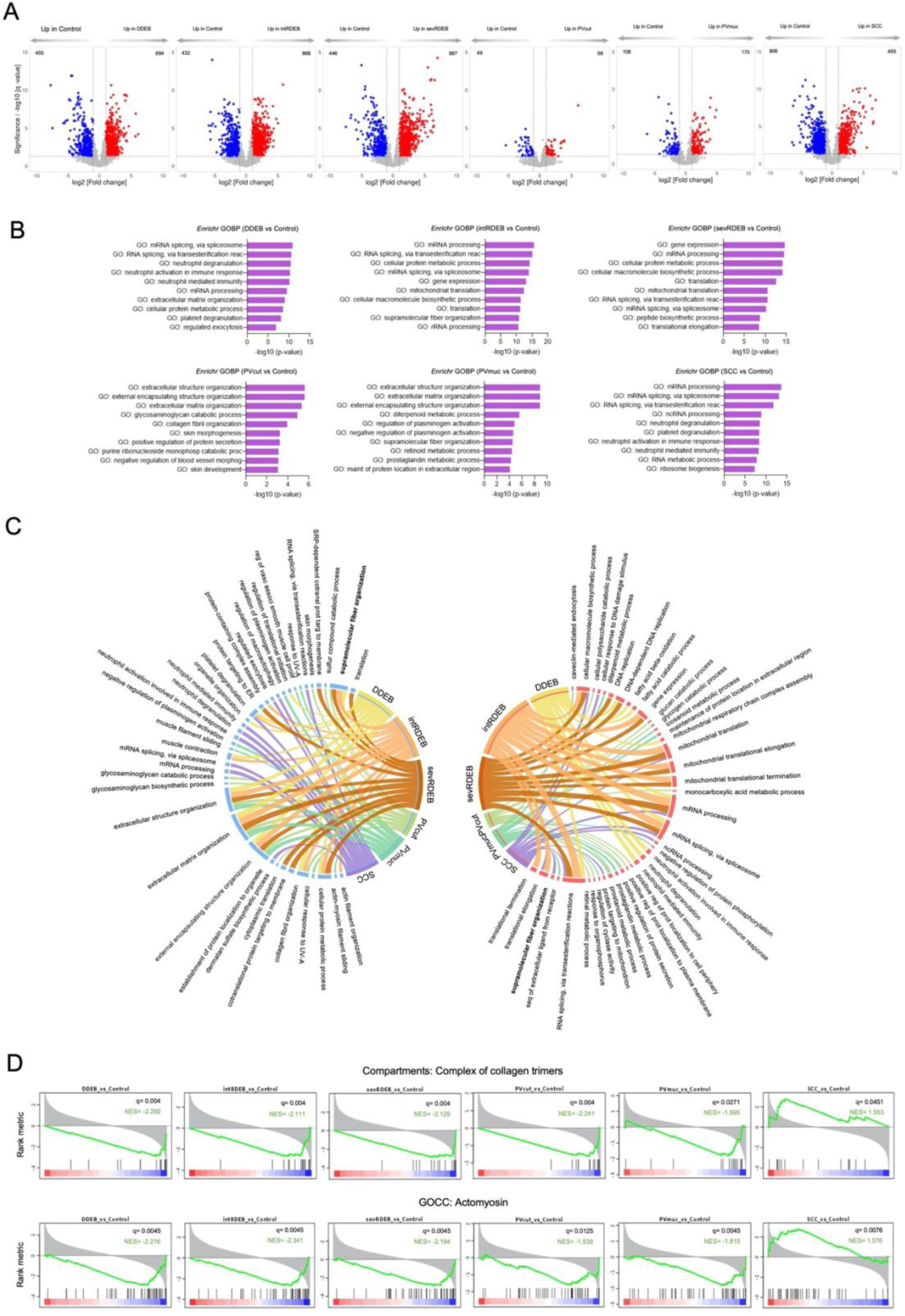
Collagen fibril and actomyosin processes are altered in all diseases but not in the same manner. **A)** Volcano plots comparing protein expression between diseases and healthy control. Differential expression was determined by the LIMMA method. Coloured values represent DEPs determined by cut-off values of fold change ± 1 and q-value < 0.05; **B)** Top 10 enriched GOBP for each comparison disease vs control. Enrichment analysis was performed using the Enrichr tool with both down- and upregulated proteins; **C)** Circos plots showing the overlap of top 10 GOPBs of downregulated (left) and upregulated (right) DEPs of each comparison disease vs control. Ribbon size represents - log10 (p-value) - wider ribbons correspond to higher enrichment; **D)** GSEA enrichment plots of the altered GO child terms under GO supramolecular fibre organization term for each comparison disease vs control. Black vertical lines represent ranked proteins in the listed signatures. Green curve represents the Normalized enrichment score (NES). The more the green line is shifted to the upper left side of the graph the more the signature is enriched in the disease. In contrast, when the green line shifts to the lower right side the higher the enrichment on the control. The red-to-blue coloured scale at the bottom represents the degree of correlation of protein’s abundance in the disease group (Red: positive; blue: negative correlation) or vice versa for the control.

We next performed enrichment analysis on DEPs from each comparison of disease versus control using the Enrichr tool [21–23]. Examination of the top 10 GOBP terms associated with each comparison revealed no common terms between them (**Figure 2B**). For DEB variants and SCC, top GOBPs were related to mRNA processing and splicing, while for PV variants top GOBPs were related to extracellular structure and matrix organization. To get further insight we did enrichment analysis for down- and up-regulated DEPs separately. We found that in the down-regulated proteins, top terms were associated with extracellular structure organization, extracellular matrix organization and external encapsulating structure organization, and were shared by all DEB and PV variants (**Figure 2C**). Of note, only one - platelet degranulation - of the top 10 GOBP terms for down-regulated DEPs in SCC was shared with two other disease variants (DDEB and intRDEB). Top GOBP terms associated with the up-regulated DEPs shared by all DEB variants and SCC were related to mRNA splicing and mRNA processing. It’s worth noting that only two of all top 10 terms for PVcut and PVmuc up-regulated DEPs were shared among them while the remaining were specific of each PV variant. Interestingly, the enrichment analysis also showed that supramolecular fiber organization GOBP was associated with SCC up-regulated DEPs and with DDEB, intRDEB, sevRDEB and PVcut down-regulated DEPs. To confirm the association of this GO term with all diseases, we carried out a pre-ranked Gene Set Enrichment Analysis (GSEA) [24–26] testing gene set signatures related to supramolecular fiber organization child terms. The only significantly enriched gene set signatures in all comparisons (diseases vs healthy control) were complex of collagen trimers and actomyosin (**Figure 2D** and **Figure S2**). Moreover, collagen fibril organization was significantly enriched only in the DEB and PVcut variants (**Figure S2**). Concomitant to the supramolecular fiber organization association to down-regulated DEPs, these three gene sets had negative normalized enrichment scores (NES) for all DEB and PV variants. In turn, SCC had a positive NES, concurrent to its up-regulated DEPs being associated with supramolecular fiber organization. The same NES pattern was also observed for Actomyosin structure organization (**Figure S2**), however without significant enrichment. In sum, GSEA showed that collagen and myosin related processes are altered in all conditions but not in the same manner.

### Collagen XII and collagen VI expression is impaired in all disease variants

Next, we investigated which proteins are potentially commonly involved in the disease alterations. We performed an intersection analysis of all down- and up-regulated proteins of each comparison of disease versus the healthy control, using a threshold of q value < 0.05. The Venn diagram intersection of each disease contrast showed that only 17 proteins were commonly significantly altered in all the diseases (**Figure 3A, B**). Interestingly, all proteins except COL3A1 and MMP2 were consistently down- or up-regulated in all diseases (**Figure 3B**). COL3A1, one of the main collagens in skin ECM, was the only common protein downregulated in DEB and PV and up-regulated in SCC, a dysregulation pattern mimicking what we observed in the Complex of collagen trimers gene set. Notably, COL12A1 was found to be significantly down-regulated in all diseases and respective variants. Since this collagen has an important role in regulating the organization of collagen fibril bundles [27–30], we performed a correlation analysis centered on COL12A1 expression to detect possible altered protein interactions (**Figure 3C**). Partial correlation network included 4 (THBS1, EDC4, HADHA and CAT) of the 17 common proteins. Additionally, other proteins related to the gene sets identified in the GSEA, such as COL5A1, MYL6 and TPM1, were also part of the network (**Figure 2D, S2**), indicating that they might also be altered but to different extents or with different expression patterns. We then looked at the expression profile of the proteins in the ontology sets complex of collagen trimers, actomyosin, collagen fibril organization and actomyosin structure organization. Additionally, we analyzed the expression of other collagens, myosins and tropomyosins (**Figure 3D**).

**Figure 3.**
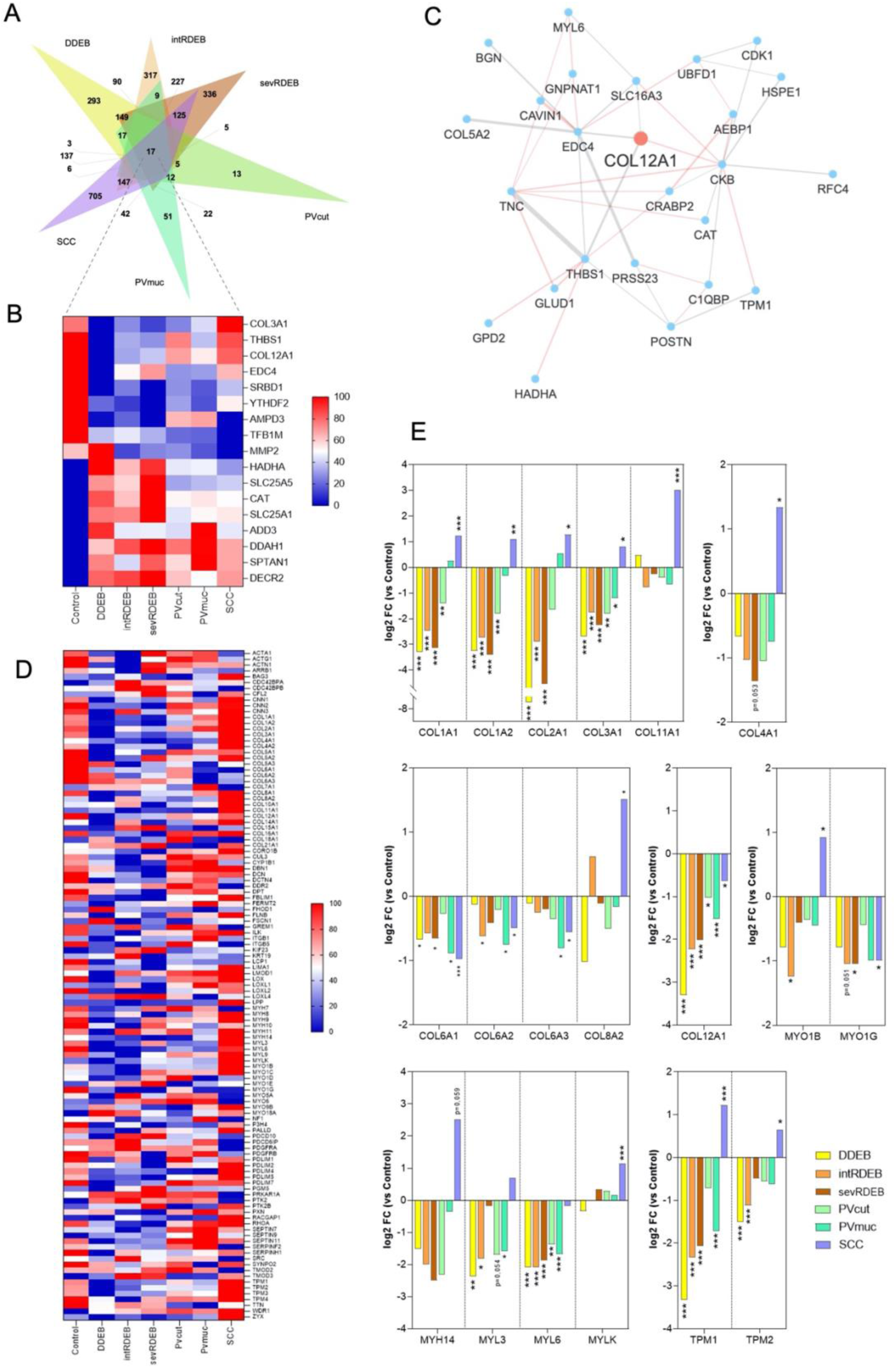
Expression of collagens, myosins and tropomysins is affected differently in disease variants. **A)** Venn diagram showing the intersection of significantly down- and up-regulated proteins (q-value < 0.05) of all comparisons of diseases versus control; **B)** Normalized heatmap of the 17 proteins commonly altered in all diseases. For each protein dark red (100) and dark blue (0) corresponds to the highest and lowest abundance, respectively, for that specific protein; **C)** Partial expression correlation network centred on COL12A1 with the top most correlated features. Red lines correspond to negative correlation and grey lines correspond to positive correlation. Width of lines is proportional to the absolute partial correlation value of the protein pair; **D)** Normalized heatmap of proteins related to the ontology sets complex of collagen trimers, actomyosin, collagen fibril organization, actomyosin structure organization together with other collagens, myosins and tropomyosins detected by LC-MS; **E)** log2 Fold Change expression (disease vs control) of selected collagens, myosins and tropomyosins. Asterisks denote (LIMMA q-value) significance against control (* q<0.05, **q<0,01, ***q<0,001).

This analysis identified that in addition to COL3A1, also COL1A1, COL1A2, COL2A1, COL11A1, MYO1B, MYH14, MYL3, MYL6, TPM1 and TPM2 had a differential expression pattern with inverse expression in SCC in relation to DEB and PV variants. However, this was not significant in all comparisons of disease vs control. Similarly to COL12A1, we observed that COL6A1, COL6A2, COL6A3 and MYOG1 were downregulated in all disease variants, although not significantly (**Figure 3E**). This is of particular interest since Collagen XII and Collagen VI defects have been implicated in the alterations of the extracellular matrix of various tissues including the skin of the Ehlers-Danlos syndrome.

### Protein expression of different skin diseases derives from shared upstream regulators

Next, we aimed at identifying if there was a common upstream regulatory environment responsible for the protein expression alterations we observed in the diseases. We used eXpression2Kinases network analysis that, for each comparison (disease vs control), computes a differential expression signature and constructs a protein-protein network to deduce upstream regulators [31]. We first performed a transcription factor (TF) analysis and compared putative enriched TFs for all diseases (**Figure 4A**). The intersection of the top 10 enriched TFs showed that there were no common deduced TFs. However, the intersection excluding PVmuc showed that MYC, MAX and GABPA were consistently among the TFs predicted to regulate the expression of DEPs of the other 5 diseases (**Figure 4B**). We also used the X2K algorithm to perform a kinase enrichment analysis feeding DEPs and predicted TFs as inputs. The results showed cyclin-dependent kinases (CDK1 and CDK4), mitogen-activated protein kinases (MAPK1/ERK2 and MAPK14/p38α) and casein kinase (CSNK2A1) consistently ranked among the top enriched kinases for all diseases (**Figure 4C, D**), showing that the alterations in all diseases potentially resulted from the activity of shared upstream regulators. Additionally, kinome tree dendrograms mapping of predicted kinases showed that these kinases belong almost exclusively to the CMGC family of kinases (including cyclin-dependent kinases – CDKs, mitogen-activated protein kinases – MAPKs, glycogen synthase kinases – GSK, and CDK-like kinases) (**Figure 4E** and **Figure S3**). Importantly, CMGC kinases and their substrates network have been shown to be involved in numerous disease processes including genetic skin disorders and skin cancers [32].

**Figure 4.**
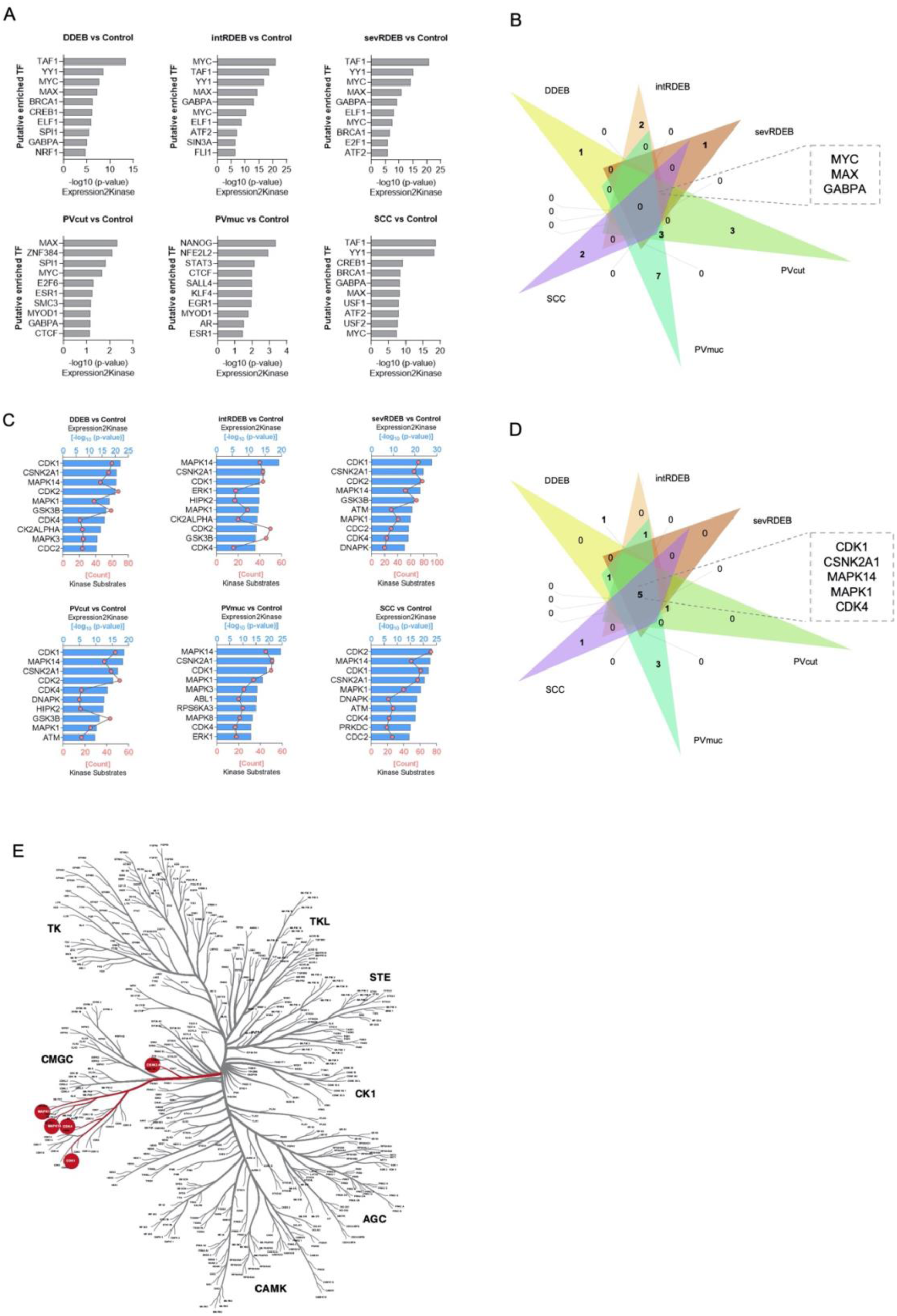
Disease alterations share an upstream regulatory environment. **A)** Bar plots showing the putative top 10 most significantly enriched transcription factors (sorted by significance level, p-value < 0,05) upstream of DEPs of each disease. Transcription factors were deduced using the transcription factor enrichment analysis module of the eXpression2Kinase web tool; **B)** Venn diagram showing the intersection of top 10 most significantly enriched transcription factors upstream of DEPs of all diseases. Common kinases to all diseases excluding PVmuc are highlighted in the box; **C)** Bar plots showing the putative top 10 most significantly kinases (sorted by significance level, p-value < 0,05) upstream of DEPs of each disease. Kinases were deduced using the Kinase enrichment analysis module of the eXpression2Kinase web tool; **D)** Venn diagram showing the intersection of top 10 most significantly enriched kinases upstream of DEPs of all disease. Common kinases are highlighted in the box; **E)** Kinome tree dendrogram mapping the common 5 kinases in the intersection of top 10 most significantly enriched kinases upstream of DEPs of all disease variants.

## DISCUSSION

Compelling evidence shows that alterations in ECM structure and composition result in dysregulation of the dermal microenvironment leading to a wide range of skin pathologies [2]. In this work we compared the proteome of fibroblasts derived from diseased skin in pursuit of common mechanistic targets with therapeutic potential. In particular, we centered our investigation in the three most prevalent DEB variants, cutaneous and mucous PV and SCC as representatives of diseases with different pathological origins.

The distinct etiology of the diseases we compared was reflected in the singularity of the protein expression maps of the disease variants, as well as in the hierarchical clustering where variants from the same disease were grouped separately from other diseases. MS-based proteomics analysis placed PV variants within the same hierarchical branch as the healthy control and revealed that PV protein expression had less differences to control than other diseases. The observed differences suggest a contribution of the dermal fibroblasts to the pathology of PV, summing to few studies that have delved into the features and role of the dermis in this disease [11,33,34]. This also aligns with recent evidence indicating that specific populations of dermal fibroblasts are responsible for the anatomic pattern of affliction in autoimmune skin disease vitiligo [3], and that dermal fibroblasts are the signal transducers between immune cells and basal cell proliferation in psoriasis [4]

The disparity in the proteome of fibroblasts across the diseases we investigated was similarly reflected in the association of DEPs to different GOBPs. However, DEB variants and SCC exhibited a common association to mRNA processing and splicing. Alterations in these processes have also been reported in studies showing that dysregulation in splicing is an important factor in the severity of DEB [35–38] and in the development of various types of SCC [39–43]. Notably, DEB patients, especially those suffering from sevRDEB, are subject to higher susceptibility to develop SCC [7,18,44]. Therefore, our results could point for a potential role for RNA splicing in the progression of SCC in DEB patients.

Moreover, our study showed that the common alterations in fibroblasts from DEB, PV and SCC diseases were only linked to the supramolecular fiber organization process. Related child terms “complex of collagen trimers” and “actomyosin” exhibited a significant enrichment in all disease versus control comparisons. Notably, the enrichment pattern observed in SCC was opposite from that seen in DEB and PV. This pattern was further reflected in protein expression, with several collagen types and myosin chains showing increased levels in SCC and decreased expression in DEB and PV. These observations are consistent with literature, as collagen synthesis and deposition is increased in many types of cancer, including SCC, pancreatic, gastric and breast cancers [45]. Increased collagen deposition due to higher actomyosin contractility has been linked to augmented tissue stiffness [46], which is also know to expedite SCC progression, including on RDEB patients [7,18]. Oppositely, inhibition of actomyosin contractility results in impaired ECM organization [47]. In this sense, in addition to the characteristic mutations in the *COL7A1* gene that lead to partial or total loss of type VII collagen in some DEB variants [2], lower expression of collagen IV and a decrease in ECM stiffness was shown in DEB dermal fibroblasts [48]. While data regarding PV is scarce, studies showed that mice injected with PV sera have increased expression of MMP9 [12], a metalloproteinase that degrades collagen I and III [49]. Motor protein non-muscle myosin II generates contractile tension in the cytoskeleton, a key process in fibroblast differentiation and migration [50]. Additionally, loss of myosin II activity results in decreased matrix stiffness [51] and cell-cell adhesion [52–54].

Interestingly, our data showed that collagen XII was the only collagen to be significantly downregulated in all disease variants. It contributes to the regulation of mechanical properties and organization of collagen fibrils [27–30], being overexpressed in fibroblast under tensile strain [55]. Several *COL12A1* mutations have been identified as the cause of myopathic Ehlers-Danlos Syndrome (mEDS), which is characterized by skin hyperextensibility and tissue fragility. Most subtypes are the result of mutations that impair the processing or the structure of collagen [56]. An alternative perspective for EDS pathogenesis [56] suggests that a defective collagen-ECM impairs the interaction between type I collagen and its receptor altering the adhesion profile of fibroblasts. This affects mechano-sensing ability and ultimately results in fibroblast dysfunction, compromised cytoskeleton dynamics, tensional homeostasis and in fragility of the tissue. Importantly, this proposal contemplates that this pathomechanism can be triggered by autoimmune or inflammatory conditions that lead to collagen aberrations in the ECM [56]. The same principle could also be applied to other diseases including those that were the focus of our study. As such, we speculate that collagen XII could be a common player mediating the pathogenic mechanism of DEB, PV and SCC. In fact, recent studies in REDB patient samples have shown alterations in ECM organization [6] and reduced collagen XII secretion [57]. Moreover, collagen fibers were found to be dispersed in SCC tumors [58] and collagen XII was also shown to mediate the progression and metastatic potential in gastric and esophageal SCC [59,60]. As for PV, to the best of our knowledge no direct targets in the dermal layer have been described, however in this study we observed the downregulation of several collagens in the dermal proteome of PVcut and PVmuc. In other immune-mediated skin diseases dermal fibroblasts were identified as responsible for the pattern of immune epidermal activity [3,4]. Therefore, there is the possibility that specific skin regions, where dermal collagen fibril assembly and organization mediated by collagen XII might fail, could be more prone to be targeted by PV autoantibodies.

Interestingly, our analysis showed that the regulatory environment responsible for the proteome alterations in each disease was also shared. MAPKs were ranked in the top 10 enriched kinases in all diseases. Several studies have highlighted the role of MAPK pathways in mediating the deposition of extracellular matrix in various cell types including chondrocytes, mesangial cells, leiomyoma cells, and keloid and dermal fibroblasts [61–67], and regulating cell adhesion and integrin expression in prostate cancer cells [68]. Strikingly, recent evidence demonstrated that MAPK signaling induced cell migration, proliferation and metastasis mechanisms are mediated by collagen XII in several cancers [59,60]. Importantly, the results of our study must be considered against a potential limitation in terms of sample size. The rarity of the diseases at study hinders patient sample availability and might limit data interpretation.

In sum, our work highlights the vast differences in the dermal proteome of skin diseases. However, we also observed a key common pattern of down regulation of collagen XII that is likely responsible for disorganization of ECM collagen fibrils and defects in cytoskeleton mechano-sensing capacity leading to skin fragility, blistering and carcinogenic phenotypes. Importantly, future work is needed to scrutinize the relationship of collagen XII with the known triggers of each of the diseases in this study and their phenotypic outcomes.

## MATERIALS AND METHODS

### Fibroblast isolation and culture

Healthy skin samples were collected from discarded tissue from adult patients who underwent abdominoplasty surgical procedures. PV and SCC skin samples were collected from patients undergoing pre-diagnostic biopsy procedures. All procedures were performed after informed consent at Hospital São João (Porto, Portugal), complying with ethical regulations regarding research involving human participants as approved by the Ethical Committee of Hospital São João (477/2020). DEB immortalized (415-EP/73/192-2013 and 415-E/2118/9-2017) hdFbs were provided by EB House Austria (MTA/2017/05/23).

Primary human dermal fibroblasts (hdFbs) were isolated from skin samples using the Whole Skin Dissociation Kit (Miltenyi Biotec) according to the manufacturer’s instructions. Cells were maintained in High Glucose Dulbecco’s modified Eagle’s medium (DMEM, Thermo Fisher Scientific) supplemented with 10% FetalClone III serum (FCIII, Hyclone), 1% L-glutamine (Thermo Fisher Scientific), and 1% antibiotic/antimycotic (Thermo Fisher Scientific), and cultured in a humidified incubator at 37°C ad 5% CO_2_. For over-confluence culture experiments hDFbs (50×10^3^/cm^2^) were cultured in medium supplemented with 50μg/mL ascorbic acid (FUJIFILM Wako Chemicals) for 14 days to promote maximum ECM deposition.

### LC-MS/MS sample preparation

Samples were lysed with lysis buffer – 100mM Tris-HCl pH 7.6, 4% sodium dodecyl sulphate (Sigma Aldrich), 100mM dithiothreitol (Sigma Aldrich, and protease inhibitor cocktail (Abcam). Lysates were homogenised with ultrasounds on ice until the buffer solution was clear (3 cycles of 2 seconds each with intervals of 1 minute). Protein concentration of the lysate was measured using the Pierce Coomassie Protein Assay Kit (Thermo Fisher Scientific). 100μg of protein sample was processed following the solid-phase-enhanced sample preparation (SP3) protocol [69] to remove all the components of the lysis buffer.

### LC-MS/MS data acquisition

Protein samples were analysed on an Ultimate 3000 liquid chromatography system coupled to a Q-Exactive Hybrid Quadrupole-Orbitrap mass spectrometer (Thermo Scientific) as described elsewhere [70]. Proteome Discoverer (version 2.5.0.400, Thermo Scientific) was used to process MS raw files. Protein identification analysis was performed with the data available in the UniProt protein sequence database for the Homo sapiens Proteome 2021_03 with 20,371 entries and a common contaminant database from MaxQuant (version 1.6.2.6, Max Planck Institute of Biochemistry). Two protein search algorithms were considered: 1) the mass spectrum library search software MSPepSearch, with the NIST huma HCD Spectrum Library (1,127,970 spectra) and 2) the Sequest HT tandem mass spectrometry peptide data base search program. An ion mass tolerance of 10 ppm for precursor ions and 0.02 Da for fragment ions was considered in both search nodes. The maximum number of allowed missing cleavage sites was set to 2. Cysteine carbamidomethylation was defined as a constant modification and peptide confidence was set to high. The Inferys rescoring node was considered for the analysis. The processing node Percolator was enabled with the following settings: maximum delta Cn 0.05; Target False Discovery Rate-FDR 1%; validation based on q-value. Protein-label-free quantification was performed with the Minora feature detector node at the processing step. Precursor ion quantification was performed at the processing step using the following settings; Peptides: unique plus razor; precursor abundance based on intensity; normalization mode was based on a t-test (background based). A chromatographic retention time alignment was applied using a maximum shift of 10 min and 10 ppm of mass tolerance allowing for mapping features from different sample files. The minimum signal-to-noise (S/N) threshold for feature linking and mapping, was set to 5.

### LC-MS/MS data analysis and visualization

#### Bioinformatics data analysis and visualization

Raw read counts were imported into *Omics Playground* (version v2.8.19, BigOmics Analytics) [71] implemented in *Docker Desktop* (Docker Inc). *Omics Playground* suite was used to perform the following analysis: Differential expression analysis was performed using the trend.LIMMA method. Gene set functional enrichment was performed using the Geneset Enrichment Analysis [24–26] *test signature* using *fgsea*. Uniform manifold approximation and projection (UMAP) [20]signature plots of protein expression were visualized using relative normalized log-expression (logCPM) and covariance as distance metric between proteins. Clustered heatmap was obtained selecting the top 500 features defined by standard deviation of protein expression. Functional annotations of heatmap clusters were defined based on geneset reference level, Fisher-weighted correlation, and the reference database Gene Ontology Biological Processes. Partial Correlation Network centered on COL12A1 was obtained using radius of 2, pcor threshold of 0.08 and Fruchterman-Reingold layout. Pairwise scatterplots were obtained by comparing differential expression profiles of all comparisons of disease versus control.

Biological processes enrichment analysis of differentially expressed proteins was performed using the Gene Ontology Biological Processes database implemented on the *Enrichr* web tool [21–23]. Analyses were performed on both upregulated and downregulated proteins, unless otherwise specified.

Volcano plots were generated using the *VolcaNoseR* web tool [72].

*GraphPad Prism* (version 9.3.1) was used to generate functional annotation and individual protein expression plots, Normalized heatmaps, and transcription factor and kinase enrichment plots.

Circos plots were generated using *Circos Table Viewer* (v0.63, Martin Krzywinski) [73] by overlapping the top 10 enriched GOBPs from upregulated or downregulated DEPs of each comparison disease versus control.

Venn diagrams were generated using the *jvenn* diagram viewer (INRA) [74].

#### Transcription factors and kinase enrichment analysis

Upstream kinases and transcription factors that are likely to regulate differentially expressed proteins identified were computationally inferred using the *eXpression2Kinases X2K Web* tool [31]. Top DEPs were uploaded into X2K Web which uses an enrichment algorithm to predict and rank probable transcription factors responsible for the regulation of the interrogated DEPs. X2K Web then builds a network of protein-protein interactions that is fed into the Kinase Enrichment Analysis feature. Predicted kinases were mapped to kinase tree dendrogram using Coral tool (Phanstiel Lab) [75] that incorporates phylogenetic relationship of kinases and allows qualitative and quantitative visualization of kinase enrichment.

#### Experimental design and statistical analysis

The number samples used in this study was 13 healthy control samples (4-5 replicates from 3 donors), 3 DDEB samples (3 replicates from 1 donor), 3 intRDEB samples (3 replicates from 1 donor), 3 sevRDEB samples (3 replicates from 1 donor), 4 PVmuc samples (4 replicates from 1 donor), 4 PVcut samples (4 replicates from 1 donor), 5 SCC samples (1 sample from 5 donors).

Statistical tests and significance levels were applied as part of the bioinformatic analysis in Omics Playground as described in the corresponding figure legends.

## Data availability

Raw data can be available on reasonable request upon publication.

## Acknowledgements

We would like to thank Hospital São João (Porto, Portugal) for providing patient samples. This work was supported by the European Research Council Consolidator Grant ERC-2016-COG-726061 (A.P.M.). Fundação para a Ciência e Tecnologia provided support with doctoral grants SFRH/BD/137766/2018 (M.D.M) and 2022.12293.BD (S.C.).

## Author contributions

L.M., M.D.M. and S.C. conducted cell experiments.

H.O. conducted LC-MS/MS experiments.

C.G-G. and T.K. provided the DEB cells.

L.M. and A.P.M. designed the study, analyzed the data and wrote the manuscript.

A.P.M. provided funding.

