## Supplementary Information for "Diseased skin dermal proteomic profiles reflect shared extracellular matrix dysregulation patterns"

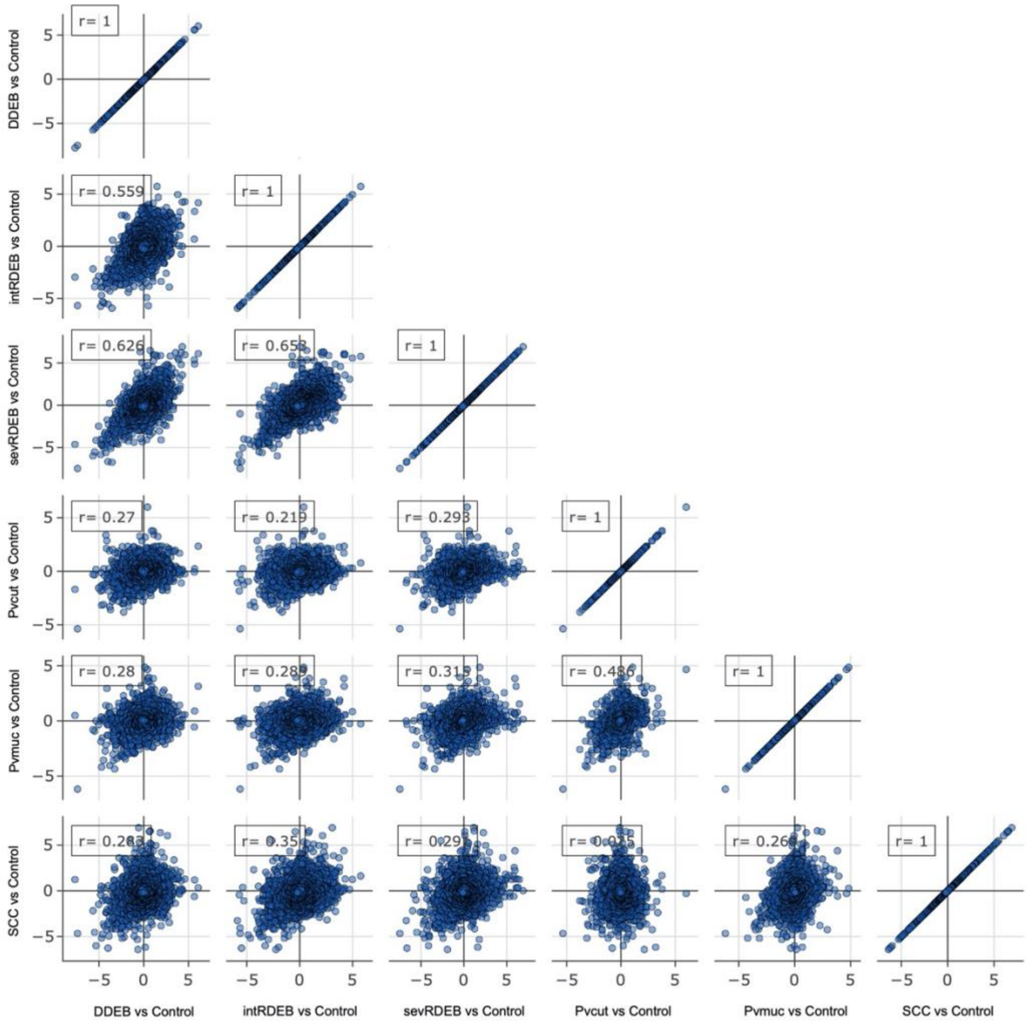


**Figure S1.** Pairwise scatterplots for the differential expression contrasts of the comparisons of disease versus control. Similar profiles show high correlation with points close to diagonal.

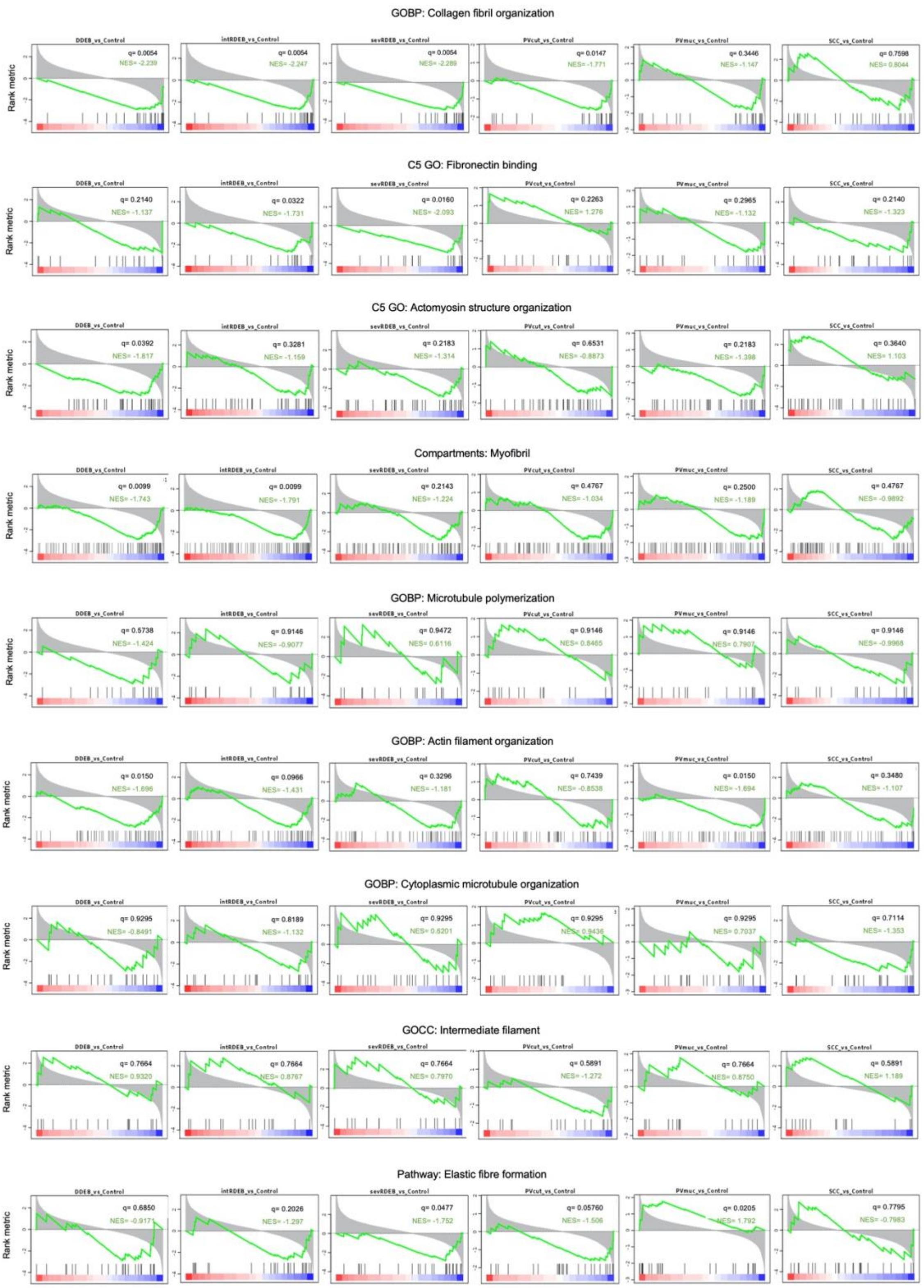


**Figure S2.** GSEA enrichment plots of the non-significantly altered GO child terms under GO supramolecular fibre organization for each comparison disease vs control. Black vertical lines represent ranked proteins in the listed signatures. Green curve represents the Normalized enrichment score (NES). The more the green line is shifted to the upper left side of the graph the more the signature is enriched in the disease. In contrast, when the green line shifts to the lower right side the higher the enrichment on the control. The red-to-blue coloured scale at the bottom represents the degree of correlation of protein's abundance in the disease group (Red: positive; blue: negative correlation) or vice versa for the control.

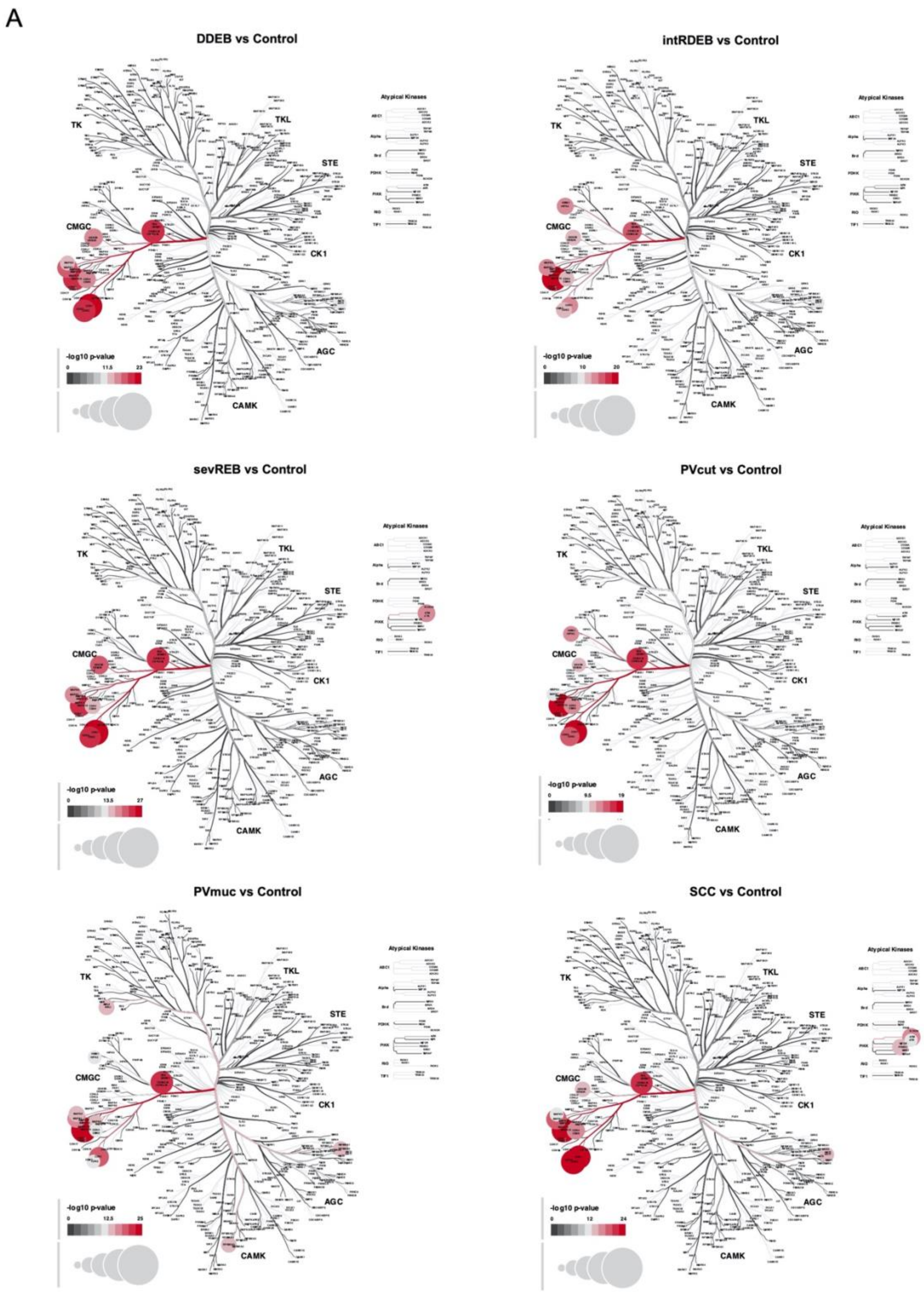


**Figure S3.** Kinome tree dendrograms mapping the enrichment of predicted upstream kinases in each disease variant. Colour and size of circles correspond to enrichment value. Dark red reflects higher p-value and dark-grey represents lower p-value.
